# Optimal Linkage Disequilibrium Splitting

**DOI:** 10.1101/2021.02.11.430793

**Authors:** Florian Privé

## Abstract

A few algorithms have been developed for splitting the genome in nearly independent blocks of linkage disequilibrium. Due to the complexity of this problem, these algorithms rely on heuristics, which makes them sub-optimal. Here we develop an optimal solution for this problem using dynamic programming. This is now implemented as function `snp_ldplit` as part of R package bigsnpr.

## Introduction

A few algorithms have been developed for splitting the genome in nearly independent blocks of linkage disequilibrium (Berisa and Pickrell 2016; Kim *et al*. 2018). Dividing the genome in multiple smaller blocks has many applications. One application is to report signals from independent regions of the genome (Berisa and Pickrell 2016; Wen *et al*. 2017; Ruderfer *et al*. 2018). Another application is for the development of statistical methods, e.g. for deriving polygenic scores (Mak *et al*. 2017; Ge *et al*. 2019; Zhou and Zhao 2020), estimating genetic architecture and performing other statistical genetics analyses (Shi *et al*. 2016; Wen *et al*. 2016). Indeed, most statistical methods based on summary statistics also use a correlation matrix (between variants), and these methods often perform computationally expensive operations such as inversion and eigen decomposition of this correlation matrix. These operations are often quadratic, cubic or even exponential with the number of variants. However, if we can decompose the correlation matrix in nearly independent blocks, then we can apply these expensive operations to smaller matrices with less variants, making these operations much faster, and parallelisable. For instance, inverting a block-diagonal matrix requires only inverting each block separately.

## Implementation

We aim at optimally splitting the genome into *K* blocks, where each block has a bounded number of variants (minimum and maximum size). This splitting is optimal in the sense that it minimizes the sum of squared correlations between variants from different blocks (hereinafter denoted as “cost”). This problem is quite complex, and a naive solution would be exponential with the number of variants. To solve this problem efficiently, we use dynamic programming, which consists in breaking a problem into sub-problems and then recursively finding the optimal solutions to the sub-problems. Here, each sub-problem consists in solving

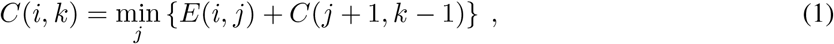

where *C*(*i*, *k*) is the minimum cost for splitting the region from variant *i* to the last variant into *k* blocks exactly, and *E*(*i*, *j*) is the error / cost between block (*i*, *j*) and the latter bloks. This is illustrated in figure 1. These sub-problems can be solved efficiently by starting with *k* = 1 and with *i* from the end of the region, and working our way up. Once all costs in the *C* matrix have been computed, and corresponding splits *j* have been recorded, the optimal split can be reconstructed from *C*(1, *K*), where *K* is the number of blocks desired. To efficiently compute 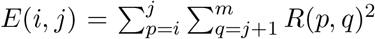, where *m* is the number of variants and *R*(*p*, *q*) is the correlation between variants *p* and *q*, we first compute the matrix *L* defined as 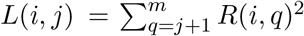. Matrices *L* and *E* are sparse. *E* is the largest matrix and requires approximately *m* × (max_size–min_size) ×4 bytes to be stored efficiently. For *m*=100,000, min_size=500 and max_size=10,000, this represents 3.5 GB.

**Figure 1:**
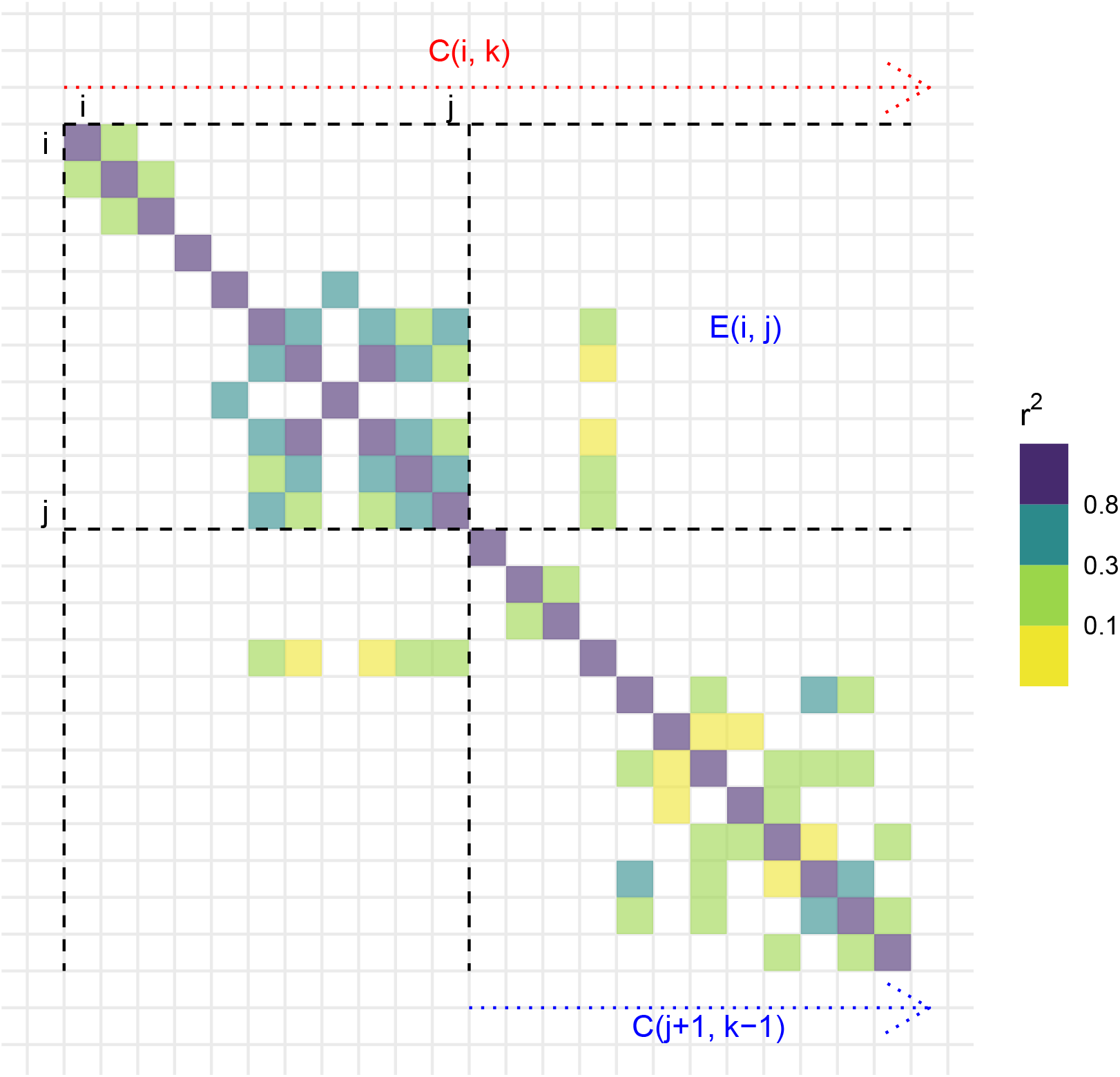
Illustration of sub-problems solved by the algorithm using a small LD matrix. The cost of separating the region starting at variant *i* in *k* blocks exactly, *C*(*i*, *k*), is broken down in two: the error *E*(*i*, *j*), the sum of all squared correlations between variants from block (*i*, *j*) and variants from all the later blocks, and the cost of separating the rest starting at (*j* + 1) using (*k* – 1) blocks. The variant *j* at which the split occurs is chosen so that the cost (*E*(*i*, *j*) + *C*(*j* + 1, *k* – 1)) is minimized.

## Results

As input, function `snp_ldsplit` uses a correlation matrix in sparse format from R package Matrix, which can be computed using the available `snp_cor` function from R package bigsnpr (Privé *et al*. 2018). This function is fast and parallelized. Then, to run `snp_ldsplit` using a correlation matrix for 102,451 variants from chromosome 1, it takes < 6 minutes on a laptop to find the optimal split in *K* blocks (for all *K* = 1 to 133) with a bounded block size between 500 and 10,000 variants. Then, the user can choose the desired number of blocks, which is a compromise between a reduced size of the blocks when using more blocks and an increased cost, since splitting a chromosome in more blocks would increase the correlation outside of these blocks. For chromosome 1 and Europeans, ldetect report 133 LD blocks (Berisa and Pickrell 2016), however we find that they can hardly be considered truly independent given the high cost (10,600) of the corresponding split (Figure S1). When splitting chromosome 1 for Europeans using the optimal algorithm we propose here, it can be split in 39 blocks at a cost of 1, in 65 blocks at a cost of 10, and in 133 blocks at a cost of 296 (Figure S1). Similar results are found for other chromosomes, and for Africans and Asians, however splitting the LD from admixed Americans comes at a high cost (Figures S2-S5)

## Software and code availability

The newest version of R package bigsnpr can be installed from GitHub (see https://github.com/privefl/bigsnpr). All code used for this paper is available at https://github.com/privefl/paper-ldsplit/tree/master/code.

