## Supplementary Materials for "Optimal Linkage Disequilibrium Splitting"

### Procedure to compute optimal costs

We use the 1000 Genomes (1000G) data in the '.bed' PLINK format provided in Privé *et al.* (2020a), and composed of 2490 individuals and 1,664,852 variants. In turn, for individuals in each of the five super populations from the 1000G data, we compute the correlation between variants with MAF $> 0.05$ using function `snp_cor` from R package bigsnpr (Privé *et al.* 2018), assuming that correlations between variants more than 3 cM away are 0 (Privé *et al.* 2020b). Then, optimal splits are computed using function `snp_ldsplit` introduced in this paper. These costs are compared with the costs provided by ldetect (Berisa and Pickrell 2016) for the European and African populations only since they are not provided for the other three super-populations. Results are presented in figures S1-S5. We see that splitting costs are very high for the admixed Americans group, showing that is impossible to split LD in independent blocks in an admixed population. This is also the case, to a lesser extent, for the African (AFR) group which includes African Americans (ASW) and African Caribbeans (ACB) who may be admixed with e.g. European ancestry. When excluding 5% of AFR individuals based on principal component analysis, splitting costs are reduced (Figure S6).

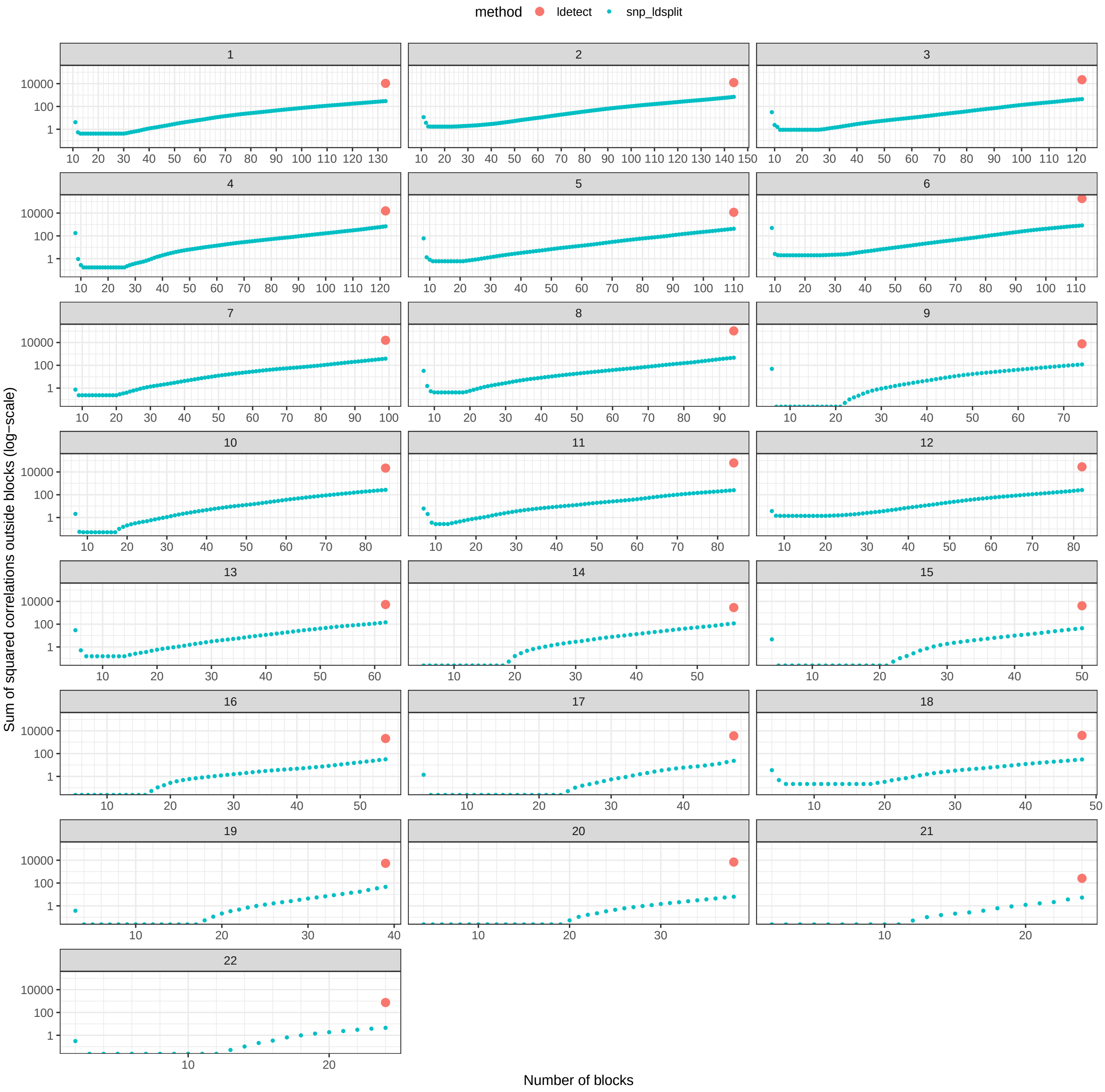


Figure S1: Optimal costs of splitting chromosomes (each panel) in a specified number of blocks (including 500 variants at minimum and 10,000 at maximum) for the European (**EUR**) super-population in the 1000 Genomes data. The cost is defined as the sum of squared correlations $r^2$ between variants from different blocks, restricting to all $r^2 > 0.05$. In red is the code from the split from ldetect, accessed from `https://bitbucket.org/nygcresearch/ldetect-data/src/master/EUR/`.

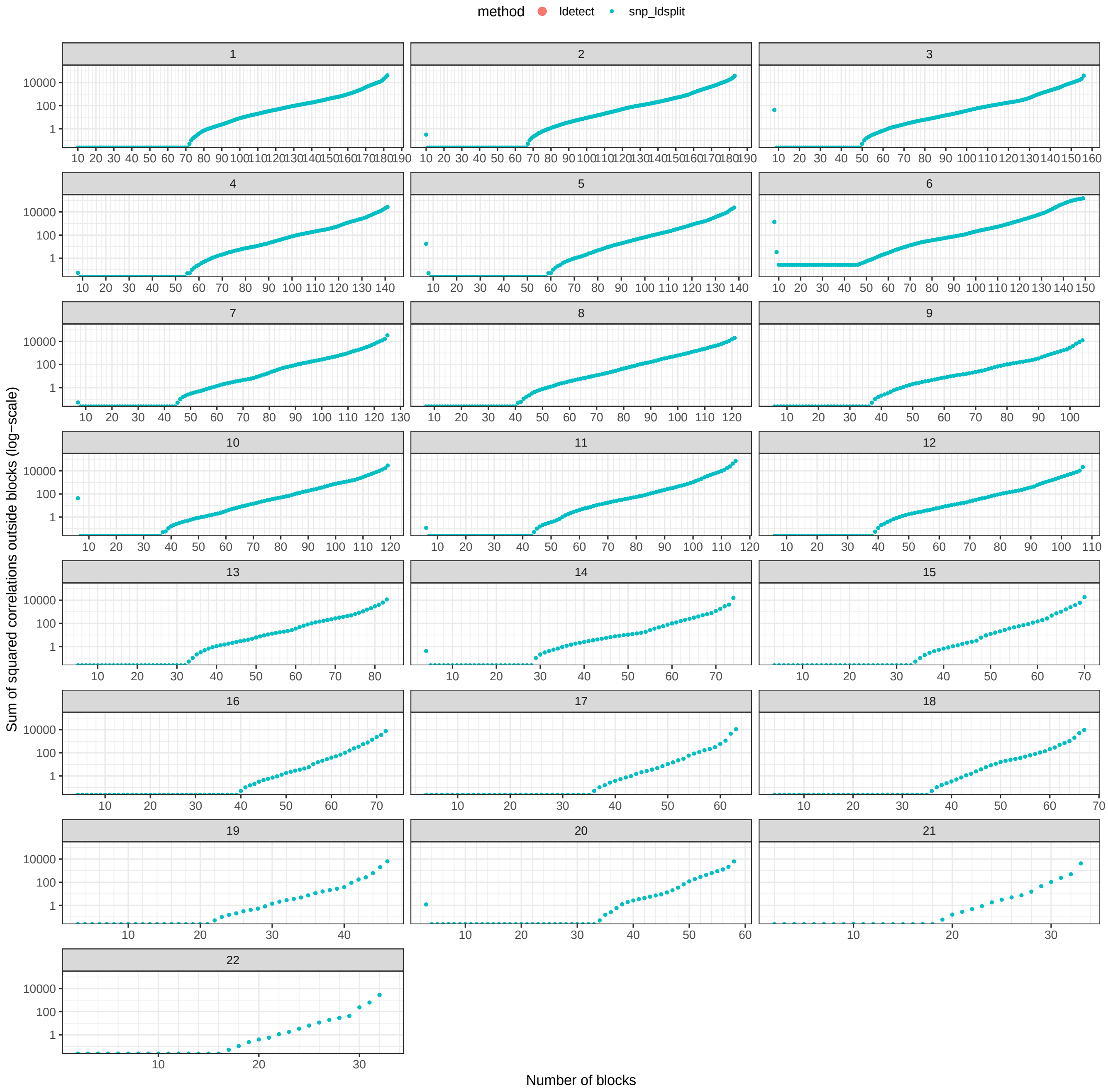


Figure S2: Optimal costs of splitting chromosomes (each panel) in a specified number of blocks (including 500 variants at minimum and 10,000 at maximum) for the East Asian (**EAS**) super-population in the 1000 Genomes data. The cost is defined as the sum of squared correlations $r^2$ between variants from different blocks, restricting to all $r^2 > 0.05$.

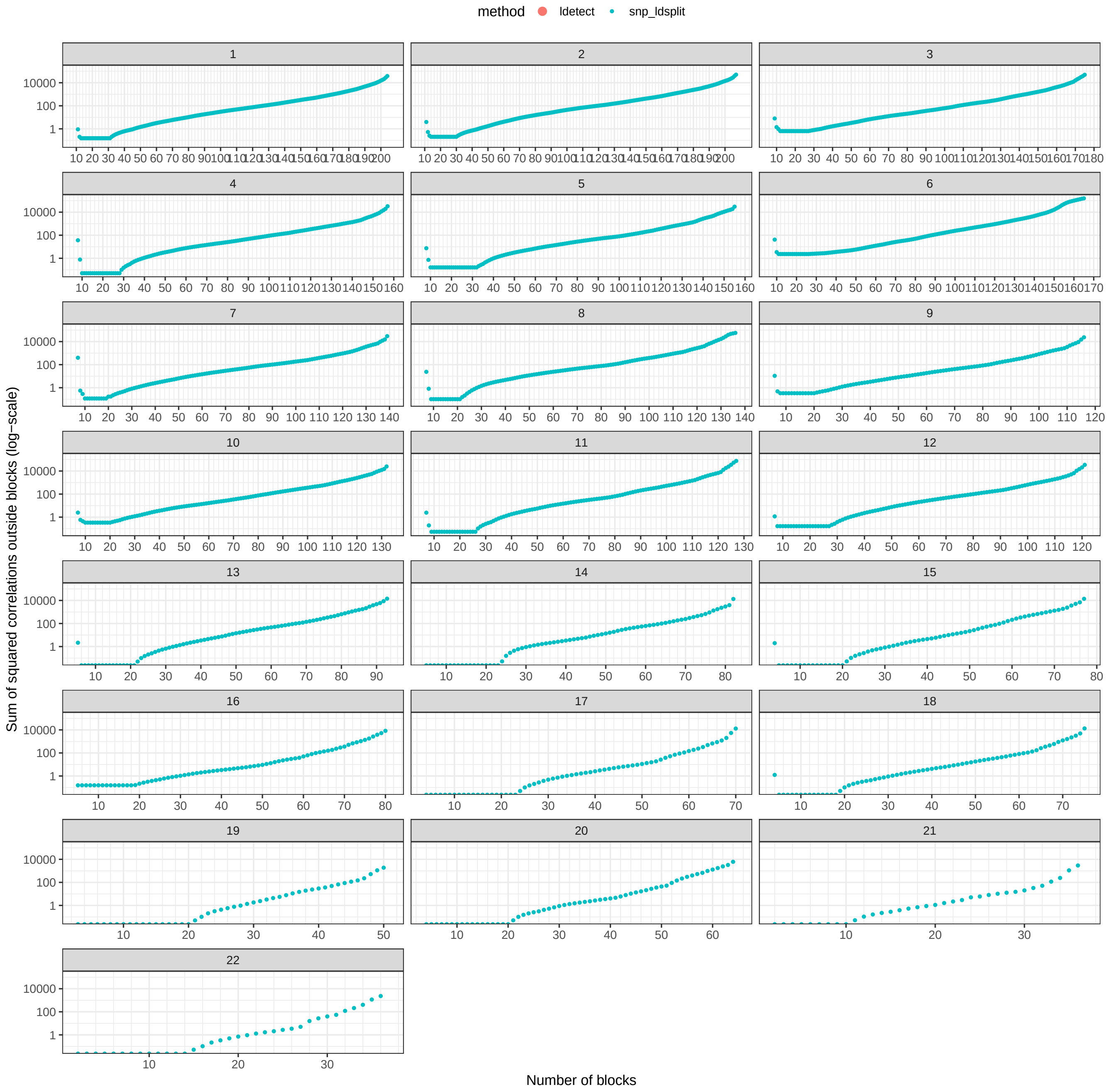


Figure S3: Optimal costs of splitting chromosomes (each panel) in a specified number of blocks (including 500 variants at minimum and 10,000 at maximum) for the South Asian (**SAS**) super-population in the 1000 Genomes data. The cost is defined as the sum of squared correlations $r^2$ between variants from different blocks, restricting to all $r^2 > 0.05$.

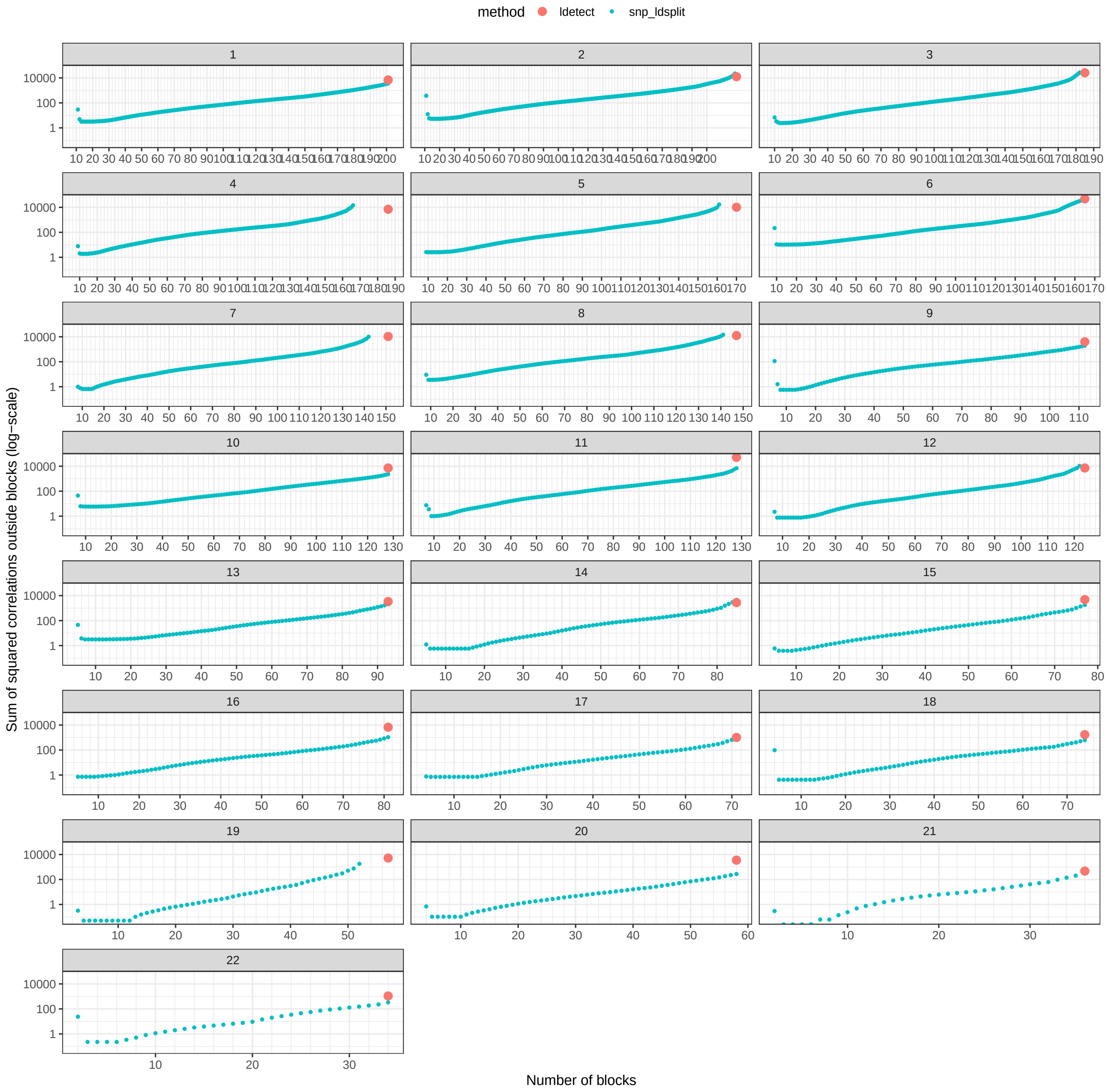


Figure S4: Optimal costs of splitting chromosomes (each panel) in a specified number of blocks (including 500 variants at minimum and 10,000 at maximum) for the African (**AFR**) super-population in the 1000 Genomes data. The cost is defined as the sum of squared correlations $r^2$ between variants from different blocks, restricting to all $r^2 > 0.05$. In red is the code from the split from ldetect, accessed from `https://bitbucket.org/nygcresearch/ldetect-data/src/master/AFR/`. Here, for the same number of blocks, the cost using ldetect can be smaller than the one with our optimal splitting since we have an additional restriction that blocks should contain at least 500 variants.

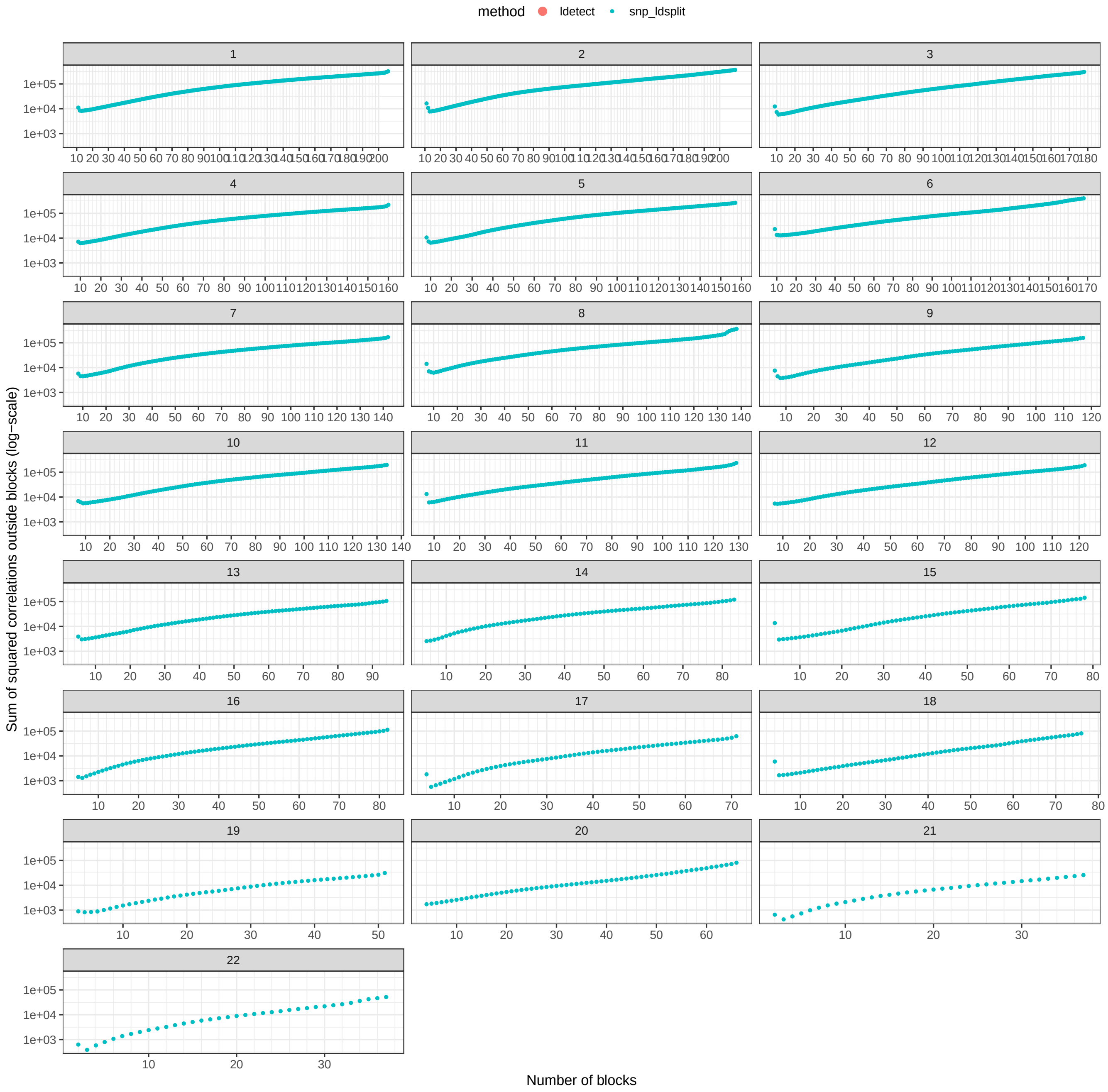


Figure S5: Optimal costs of splitting chromosomes (each panel) in a specified number of blocks (including 500 variants at minimum and 10,000 at maximum) for the Admixed American (**AMR**) super-population in the 1000 Genomes data. The cost is defined as the sum of squared correlations $r^2$ between variants from different blocks, restricting to all $r^2 > 0.05$.

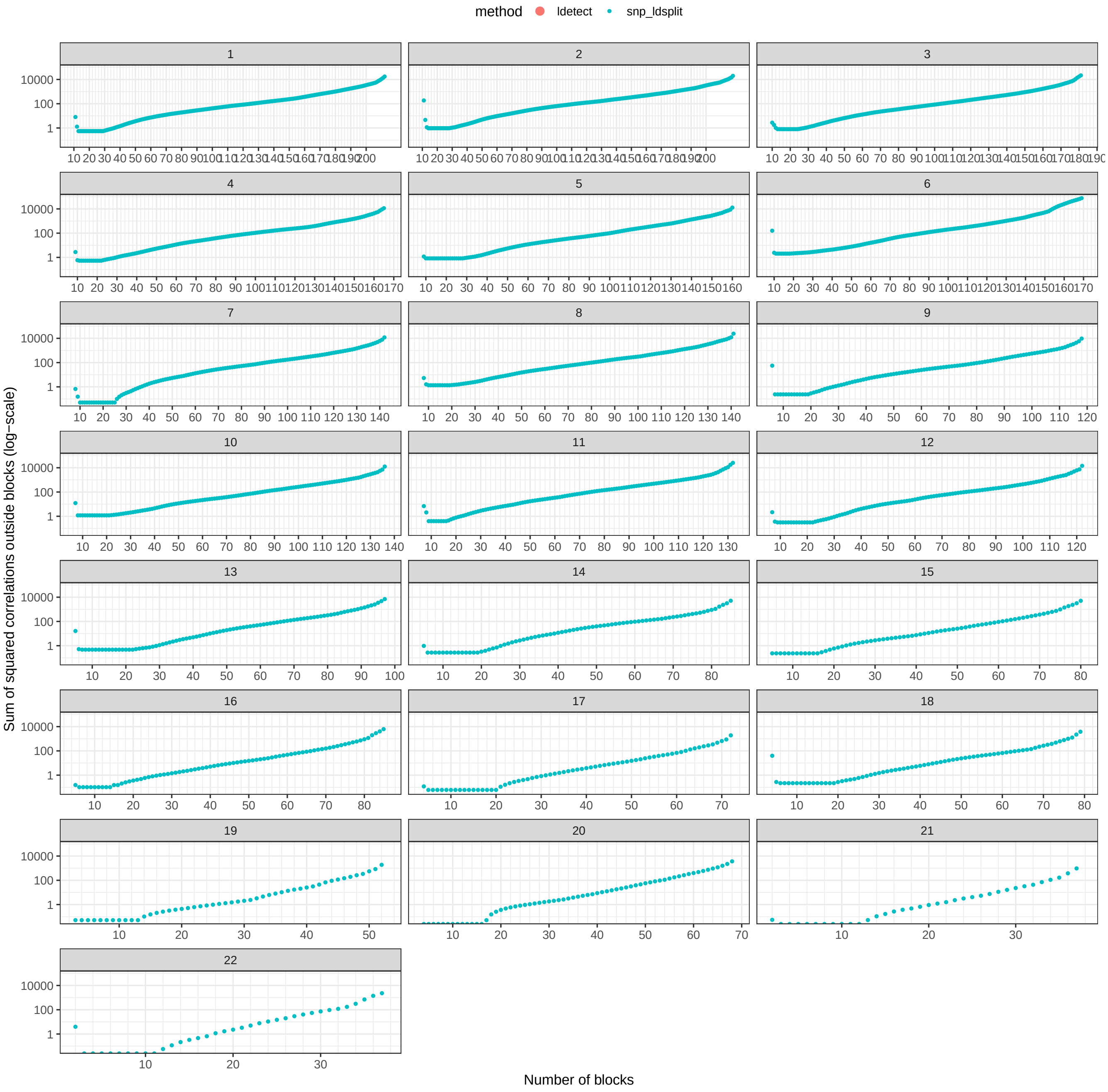


Figure S6: Same as figure S4, but after excluding 5% of AFR individuals based on PCA.
